# Nanopore-only assemblies for genomic surveillance of the global priority drug-resistant pathogen, *Klebsiella pneumoniae*

**DOI:** 10.1101/2022.06.30.498322

**Authors:** Ebenezer Foster-Nyarko, Hugh Cottingham, Ryan R. Wick, Louise M. Judd, Margaret M. C. Lam, Kelly L. Wyres, Thomas D. Stanton, Kara K. Tsang, Sophia David, David M. Aanensen, Sylvain Brisse, Kathryn E. Holt

## Abstract

**Background:** Oxford Nanopore Technologies (ONT) sequencing has rich potential for genomic epidemiology and public health investigations of bacterial pathogens, particularly in low-resource settings and at the point of care, due to its portability and affordability. However, low base-call accuracy has limited the reliability of ONT data for critical tasks such as antimicrobial resistance (AMR) and virulence gene detection and typing, serotype prediction and cluster identification. Thus, Illumina sequencing remains the standard for genomic surveillance despite higher capital and running costs.

**Methods:** We tested the accuracy of ONT-only assemblies for common applied bacterial genomics tasks (genotyping and cluster detection, implemented via Kleborate, Kaptive and Pathogenwatch), using data from 54 unique Klebsiella pneumoniae isolates. ONT reads generated via MinION with R9.4 flowcells were basecalled using three alternative models (Fast, High-accuracy (HAC) and Super-accuracy (SUP), available within ONT’s Guppy software), assembled with Flye and polished using Medaka. Accuracy of typing using ONT-only assemblies was compared with that of Illumina-only and hybrid ONT+Illumina assemblies, constructed from the same isolates as reference standards.

**Results:** The most resource-intensive ONT-assembly approach (SUP basecalling, with or without Medaka polishing) performed best, yielding reliable capsule (K) type calls for all strains (100% exact or best matching locus), reliable multi-locus sequence type (MLST) assignment (98.3% exact match or single-locus variants), and good detection of acquired AMR genes and mutations (88% – 100% correct identification across the various drug classes). Distance-based trees generated from SUP+Medaka assemblies accurately reflected overall genetic relationships between isolates; however, the definition of outbreak clusters from ONT-only assemblies was problematic. HAC basecalling + Medaka polishing performed similarly to SUP basecalling without polishing, and polishing introduced errors into HAC- or Fast-basecalled assemblies. Therefore, we recommend investing compute resources into basecalling (SUP model) over polishing, where compute resources and/or time are limiting.

**Conclusions:** Overall, our results show that MLST, K type and AMR determinants can be reliably identified with ONT-only data. However, cluster detection remains challenging with this technology.

## Introduction

*Klebsiella pneumoniae* is a highly versatile species widely linked with multi-drug resistant (MDR) healthcare-associated infections and hypervirulent community-acquired infections [1, 2]. It contributes a third of all healthcare-associated Gram-negative infections globally and is a leading cause of Gram-negative invasive diseases, second to *Escherichia coli* [1–3]. Worryingly, the acquisition rate of antimicrobial resistance (AMR) determinants among *K. pneumoniae* has continued to rise rapidly over the last decades—particularly resistance to third-generation cephalosporins and carbapenems. The Global Research on AntiMicrobial resistance (GRAM) study estimated that *K. pneumoniae* was one of six pathogens that caused more than 250,000 deaths associated with AMR in 2019 [4]. In that study, *K. pneumoniae* was the leading Gram-negative pathogen in sub-Saharan Africa, contributing 20% of the deaths attributable to AMR [4]. At the same time, data aggregated from 151 studies encompassing 26 countries in sub-Saharan Africa indicate that *K. pneumoniae* is the top Gram-negative and second most important cause of neonatal sepsis [3]. *K. pneumoniae* infections are also characterised by a high case fatality rate (18-49%) [1, 3, 5, 6]. Consequently, *K. pneumoniae* has been flagged in the World Health Organisation’s highest category of priority pathogens, for which the development of new antibiotics is urgently needed [7].

Accumulating genomic evidence has highlighted the emergence of hybrid MDR-hypervirulent clones, driven by the acquisition of mobile genetic elements [5, 6, 8–10]. This phenomenon represents a significant challenge to treating infections caused by *K. pneumoniae* and signals the urgent need for appropriate surveillance systems to identify MDR-hypervirulent clones and their genetic elements. WHO recommends the robust surveillance of AMR as a critical part of the Global Action Plan on AMR [11]. However, classical approaches to surveillance have relied on antimicrobial susceptibility testing and low resolution genotyping methods, which yield little information on the evolution and spread of AMR within bacterial populations.

AMR surveillance in critical pathogens such as *K. pneumoniae* has been dramatically enhanced by the recent democratisation of whole-genome sequencing (WGS), primarily by short-read (Illumina) sequencing. WGS is currently the gold standard approach for public health pathogen surveillance, facilitating cluster identification and genotyping of clinically relevant loci, including AMR, virulence traits, and antigen prediction [12–15]. Furthermore, as few labs perform *K. pneumoniae* serology, prediction of capsular (K) and lipopolysaccharide (O) antigen serotypes and their population distribution relies on inference from K and O antigen biosynthesis loci captured in whole-genome sequence data [16, 17]. Genomic surveillance for K and O types is of increasing interest, as these antigens are the main target for alternative control measures such as vaccines, phage therapy, and monoclonal antibody therapy [18–21]. Thirteen O antigen locus (OL) and 162 K antigen locus (KL) types are described to date [16]. Unfortunately, the gold standard Illumina short-read sequencing platforms result in fragmented genome assemblies, which cannot accurately resolve plasmids and other mobile genetic elements that drive the dissemination of AMR determinants via horizontal and lateral gene transfer. Fragmentation of Illumina genome assemblies also complicates K and O typing, resulting in up to a third of Illumina-based genomes being untypeable [22]. Long-read sequencing platforms, such as Pacific Biosciences and Oxford Nanopore Technologies (ONT), can overcome these limitations, as they yield longer reads (often exceeding 10 kbp, with current maximal read lengths reaching 1 Mbp) that are capable of resolving structural variations, long repeat regions, and genomic copy-number alterations [23–25]. ONT is increasingly adopted to generate high-quality hybrid Illumina-plus-long-read assemblies for bacteria, owing to its flexibility and low capital cost compared to PacBio [26, 27]; however, high read-level error rates have precluded precluded the adoption of ONT devices as a standalone platform for pathogen genomic epidemiology studies or public health investigations [28]. The situation has shifted somewhat during the COVID-19 pandemic as ONT has been widely adopted for virus sequencing and outbreak analysis, including in settings where Illumina platforms are not available and are unlikely to become so due to capital costs, reagent access, and other logistical issues [29]. There is, therefore, now an unparalleled opportunity to harness ONT sequencing for genomic epidemiology and surveillance of bacterial pathogens, which is particularly attractive for AMR-associated pathogens such as *K. pneumoniae,* where resolving plasmids and other highly variable genomic components is highly beneficial.

The current state-of-the-art ONT basecaller, Guppy, utilises the Fast, High-accuracy (HAC) and Super-accuracy (SUP) models. The Fast model, as the name indicates, is the fastest of the three algorithms but at the cost of accuracy. SUP is the most accurate and slowest; the speed of the basecaller being a factor of the number of parameters in the neural network model. HAC is intermediate between Fast and SUP and about six times slower than the Fast model (https://github.com/rrwick/August-2019-consensus-accuracy-update#basecalling). ONT polishing, utilising Medaka—a neural network-based tool that generates consensus sequences by aligning the individual sequence reads against a draft assembly [30]—is rapid, highly effective and believed to improve sequence accuracy significantly [31]. ONT continuously refines these algorithms to improve raw reads and consensus accuracy [32].

However, despite continuous improvement over the years, ONT is still susceptible to non-trivial numbers of base-call errors, which can impact the identification of clinically important features such as acquired virulence and AMR genes [33, 34]. For example, a mean global error rate of ∼6% is reported for reads with quality scores of at least 10—corresponding to 94% raw read accuracy and >99.99% consensus accuracy with the primary nanopore in use currently (R9.4.1 chemistry), although this varies by the organism [35] and is improving with new chemistries such as R10 [36]. For a 5 Mbp genome, such as *K. pneumoniae,* we previously reported roughly 3,000 errors (corresponding to at least 337 substitutions per genome assembly) [37]; therefore, ONT-only assemblies are prone to erroneous SNP calls, which can potentially hamper outbreak investigations [38]. The performance of ONT-only assemblies in identifying AMR and virulence traits against the gold-standard (hybrid assemblies) is unclear but is gaining attention [39, 40], and is pertinent to inform the research community working with long-read data on what levels of accuracy to expect with current basecalling algorithms.

Here, we use a collection of 54 clinical isolates of *K. pneumoniae* with matched Illumina and ONT data [41] to assess the performance of ONT-only data when used for the prediction of AMR determinants, virulence factors, K/O typing and phylogenetic clustering analysis. Current read-based genotyping tools for Illumina data (such as ARIBA [42] or SRST2 [43]) are not optimised to be used efficiently on noisy long reads like those produced by ONT. However, assembly-based tools such as Pathogenwatch [44], Kleborate [17, 45] and Kaptive [16] can in principle be applied to any assembly regardless of the sequencing platform, allowing for direct comparisons and facilitating the straightforward application of common genotyping workflows to data generated from different sequencing instruments deployed for pathogen surveillance. Furthermore, Pathogenwatch provides a free and accessible online analysis platform, suitable for non-informatics specialists such as public health or clinical microbiologists and epidemiologists, which, together with the growing network of ONT-equipped labs could facilitate the widespread acceleration of genomic surveillance for AMR pathogens. We therefore focused our investigation on the performance of these tools (Pathogenwatch, Kleborate and Kaptive) for genomic surveillance of *K. pneumoniae* using ONT-only assemblies.

## Methods

### Bacterial isolates and sequencing

We utilised 54 *K. pneumoniae* that were previously isolated at the Alfred Health Clinical Microbiological Diagnostic Laboratory in Melbourne, Australia, as part of a year-long prospective study of isolates from clinical infections (hospital-wide) and screening swabs (in intensive care and geriatric wards) in the Alfred Health network [41, 46–48]. The accession numbers for all genome data analysed in this study are provided in **Supplementary Table 1**.

Isolates were sequenced first via Illumina HiSeq 2500, generating 125 bp paired-end reads, as described previously [46]. Later, selected isolates were subcultured, and fresh DNA extracted for long-read sequencing using GenFind (Beckman Coulter) kits (full protocol available under DOI: 10.17504/protocols.io.p5mdq46). ONT libraries were prepared using the ligation protocol with barcoding kits to multiplex 12 to 24 isolates per run (barcode kits EXP-NBD104 and EXP-NBD114) and sequenced on a MinION device with R9.4.1 flowcells as previously described [26].

### Basecalling and assembly

Fast5 files generated by the ONT MinION were initially basecalled with Guppy v4.0.14 [49] on the Monash University M3 MASSIVE GPU cluster using the “Fast” (r9.4.1_450bps_fast) and “High-accuracy”, aka HAC (r9.4.1_450bps_hac) models for each sample. The “Super-accuracy” (r9.4.1_450bps_sup) model was later used when it was released in Guppy v5.0.7 [49, 50]. The basecalled FASTQ files were then concatenated into a single file per basecalled sequence run and demultiplexed into individual FASTQ files (one per sample) with the qcat command-line tool v1.1.0 [51] based on the barcode sequences. As a quality control step before assembly, we first filtered out poor-quality reads using fastp v0.20.1 [52] (short reads) and Filtlong v0.2.1 [53] (long reads) at a sequence similarity threshold of 90-95%, retaining only the reads with a minimum length of 1 kbp and excluding the worst 5-10% of the reads (for FASTQ files less than 200 MB, a keep_percent of 95% was used, while a keep_percent of 90% was applied to read files of size greater than 200 MB).

Three separate approaches were employed to generate genome assemblies (**Figure 1**). First, Unicycler v0.4.9b [54] was used to create Illumina-only assemblies for all the study isolates using default settings. When given short reads, Unicycler performs genome assembly using SPAdes v3.14.0 [55] but includes extra fine-tuning steps such as filtering out low-depth contigs and thus low-level contamination.

**Figure 1.**
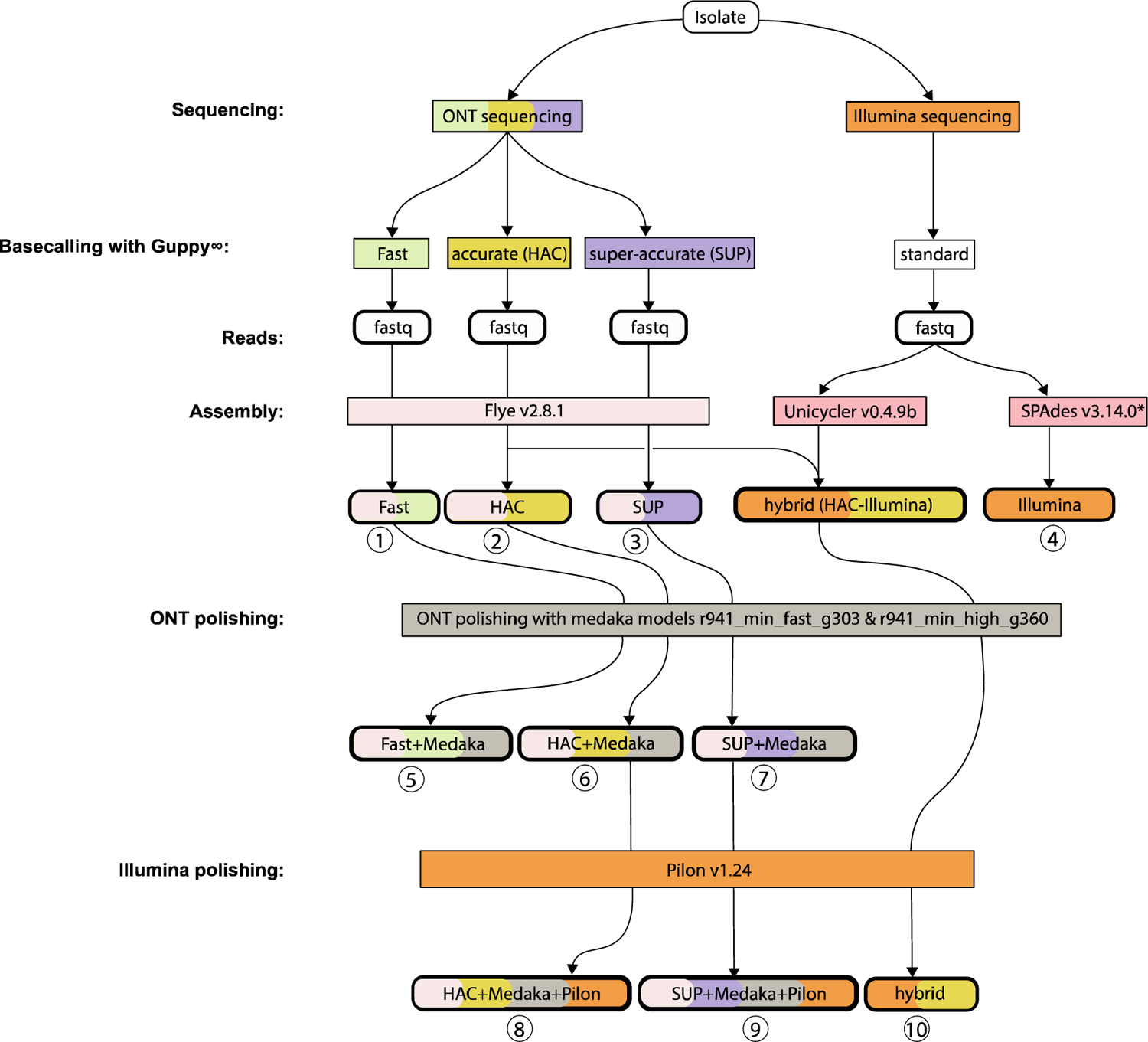
Genome assembly flow diagram. We generated ten assemblies for each of 54 *Klebsiella pneumoniae* isolates, which were sequenced on both Illumina and ONT MinION (numbered 1 to 10). ONT reads were basecalled using three alternative models (via Guppy v4.0.14): fast, high-accuracy (HAC), and super-accuracy. The resulting basecalled read sets were each subjected to assembly using Flye (v2.8.1), generating three ONT-only assemblies per isolate. These assemblies were each then polished with the corresponding ONT read set (using Medaka models r941_min_fast_g303 and r941_min_high_g360) to generate a further three ONT-only polished assemblies per isolate, totalling 6 ONT-only assemblies (3 polished, 3 unpolished) per isolate. For comparison, we used Illumina data to generate Illumina-only and ONT/Illumina hybrid assemblies for each isolate. Illumina reads were assembled using SPAdes v3.14.0 to generate an Illumina-only assembly. A short-read-first hybrid assembly was generated using Unicycler v0.4.9b and then polished using Pilon v1.24. Finally, two long-read-first hybrid assemblies were generated by polishing the ONT-only HAC and super-accuracy models using Illumina reads (via pilon v1.24). *SPAdes was run via Unicycler v0.4.9b. ∞The Fast and HAC basecalling utilised Guppy v4.0.14 while SUP basecalling was done using Guppy v5.0.7.

Next, we used Flye v2.8.1 [56] to generate ONT-only assemblies from basecalled reads (three read sets per genome, called with the three different basecallers). Flye was chosen based on our previous benchmarking of algorithms for ONT-only assemblies of bacterial genomes [57]. Each ONT assembly was then polished with ONT reads using Medaka (v1.4.3) models r941_min_fast_g303 and r941_min_high_g360 [30], resulting in two ONT-only assemblies per basecalling model (polished vs unpolished); i.e., a total of six ONT-only assemblies per sample.

Finally, we produced two kinds of hybrid Illumina+ONT assemblies: short-read-first and long-read-first. Short-read-first hybrid assemblies were generated using Unicycler v0.4.9b, which starts by building a short-read assembly graph with SPAdes v3.14.0, then uses the corresponding long reads (HAC basecalled, in this case) to scaffold the genome, and finally runs Pilon v1.23 [58] in an attempt to fill gaps, correct bases and fix misassemblies using the short reads [26, 54] (HAC basecalled reads were used for the short-read-first assemblies as these were generated before the SUP basecalling model became available [50]). To generate long-read-first assemblies [59], we used Flye v2.8-1 to produce ONT-only assemblies for the set of reads basecalled with the HAC and SUP-accuracy basecalling models, followed by long-read polishing with Medaka [30] to repair any residual errors using ONT long reads, then finally short-read polishing using Illumina reads and Pilon v1.24 (following the recommendations noted at [60]). Thus, altogether, we produced ten assemblies per sample, encompassing reads derived from the three separate basecalling models (**Figure 1**). QUAST v5.0.0 [de6973bb] [61] and CheckM v1.1.10 [62] were used to assess the quality and completion of the hybrid assemblies (See **Supplementary File 1**, available at https://figshare.com/s/6026855223031e769d8a, (DOI: 10.6084/m9.figshare.19745608).

We designate the ONT-only assemblies, with and without Medaka ONT-read polishing, as “Fast”, “Fast+Medaka”, “HAC”, “HAC+Medaka”, “SUP”, “SUP+Medaka”. Based on the recommendations of Wick et al. ([60] and [63]), we consider the long-read-first hybrid assembly derived from SUP basecalled reads as the most accurate assembly and designate this as the reference sequence, against which the performance of other assemblies is benchmarked. We hereafter use the term “hybrid” to refer to the short-read-first (Unicycler) hybrid assemblies, to distinguish them from long-read-first hybrid assemblies (otherwise referred to as “SUP+Medaka+pilon” and “HAC+Medaka+pilon”).

### Genotyping

Genome assemblies were uploaded to Pathogenwatch v2.3.1 [44] where Kleborate v2.2 [45] and Kaptive v2.0 [16] were automatically deployed to call multi-locus sequence types (STs) using the 7-locus scheme [64], capsular polysaccharide (K) and lipopolysaccharide O locus types and serotype predictions, acquired virulence traits including the siderophores aerobactin (*iuc*), yersiniabactin (*ybt*) and salmochelin (*iro*), the genotoxin colibactin (*clb*) and the hypermucoidy locus (*rmpADC*). Pathogenwatch also deploys Kleborate to identify established AMR determinants (acquired genes and chromosomal mutations) [45] for the following antimicrobial classes: aminoglycosides, carbapenems, third-generation cephalosporins, third-generation cephalosporins plus β-lactamase inhibitors, colistin, fluoroquinolones, fosfomycin, penicillins, penicillins + β-lactamase inhibitors, phenicols, sulfonamides, tetracyclines, tigecycline and trimethoprim.

### Clustering

We downloaded the Pathogenwatch pairwise distance matrix and corresponding neighbour-joining tree for the full set of assemblies. The distance matrix is available at https://figshare.com/s/6026855223031e769d8a (DOI: 10.6084/m9.figshare.19745608), and the tree for interactive viewing at Microreact (https://microreact.org/project/sUrpBsvXi1aiKD7ssPv9pu-nanopore-only-assemblies-for-genomic-surveillance-of-klebsiella-pneumoniae). Pathogenwatch calculates pairwise SNP distances between genomes based on a concatenated alignment of 1,972 genes (2,172,367bp) that make up the core gene library for *K. pneumoniae* in Pathogenwatch and infers a neighbour-joining tree from the resulting pairwise distance matrix [44]. Here, we assessed the feasibility of identifying potential nosocomial transmission clusters using these distance matrices. Several studies have proposed thresholds in the range of 21-25 genome-wide SNPs for identifying nosocomial transmission clusters of *K. pneumoniae* [65–67]. However, as Pathogenwatch calls SNPs only in 1,972 core genes and not genome-wide, we compared the SNP distances calculated by Pathogenwatch with genome-wide SNP counts obtained by mapping short reads to a reference genome to determine the equivalent cut-off for clustering analysis using Pathogenwatch distances. To do this, we used the genome-wide SNP alignment generated previously for n=270 *K. pneumoniae* isolated at Alfred Health, based on mapping of Illumina reads to the *K. pneumoniae* NTUH-K2044 reference genome using the RedDog pipeline [68] (see full details in [41]). Pairwise SNP counts were extracted using snp-dist [69]. Assemblies for these 270 genomes (assembled from Illumina reads *de novo* using SPAdes optimised with Unicycler v0.4.74, see full details in [41]) were uploaded to Pathogenwatch, and the pairwise distance matrix was downloaded and compared against that generated from RedDog. We then used R to fit a linear regression model for Pathogenwatch distances as a function of genome-wide mapping-based SNP distances (see **Supplementary Figure 1**). This indicated a Pathogenwatch distance threshold of 10 SNPs would be approximately equivalent to the established genome-wide distance threshold of 25. These thresholds assume accurate basecalling from Illumina data. To ascertain a corresponding threshold distance using ONT-only data, we compared pairwise Pathogenwatch distances calculated using ONT-only (SUP+Medaka) assemblies vs Illumina assemblies, for pairs of strains linked via probable transmission clusters. Using R to fit a linear regression model indicated an ONT-only Pathogenwatch distance of 50 SNPs would approximate the Illumina-based Pathogenwatch distance of 10 SNPs or genome-wide distance of 25 SNPs (see **Figure 6**).

We compared the topologies of neighbour-joining trees generated from Pathogenwatch distance matrices calculated using SUP+Medaka, HAC+Medaka or Fast+Medaka assemblies against the reference tree (calculated from hybrid SUP+Medaka+pilon assemblies), using the tanglegram function in the R package dendextend v1.15.2 to generate comparative tree plots and calculate entanglement coefficients. We also used the phytools package v1.0-3 in R to compute the Robinson-Foulds distance [70, 71] between tree topologies, which represents a sum of the number of partitions inferred by the first tree but not the second tree and that inferred by the second tree but not the first tree.

### Data analysis and visualisation

Data were analysed and visualised using R v4.1.0. Linear regression was done using the lm function in base R, forcing the line through the origin. Genotyping and clustering results were visualised and compared using R packages ape v5.5 [72], cowplot v1.1.1 [73], data.table v1.14.2 [74], DECIPHER v2.20.0 [75], dendextend v1.15.2 [76], dplyr v1.0.7 [77], ggtree v3.0.4 [78], ggpubr v0.4.0 [79], gridExtra v2.3 [80], janitor 2.1.0 [81], kableExtra v1.3.4 [82], phytools v1.0.3 [71], tidyverse v1.3.1 [83], treeio v1.16.2 [84], treemap v2.4-3 [85] and stringr v1.4.0 [86].

## Results and Discussion

We investigated the accuracy of ONT-only assemblies for identifying ST, K and O antigen loci, AMR determinants, plasmid- and ICE*Kp-*borne virulence factors and phylogenetic clusters, compared with assemblies generated via Illumina or hybrid methods that combine Illumina and ONT reads (**Figure 1**). We used a diverse set of 54 unique *K. pneumoniae* isolates (see **Methods**), spanning thirty STs and displaying a wide range of K types, O types, AMR and virulence profiles (see **Supplementary Table 1**). The genome sizes were in the range 5,075,945 bp – 6,163,371 bp, and spanned G+C content of 49.6% to 57.7% (**Supplementary Figure 2**). Notably, the choice of ONT basecalling model did not appear to have a significant impact on the N50 or G+C content. Full details of sequence quality and assembly metrics are presented in **Supplementary Table 1** and **Supplementary File 1**.

### MLST and clustering analysis

Perfect ST calls require exact sequence matches at all 7 MLST loci; however, ST is often treated as a proxy for lineage, which can sometimes be correctly identified based on single-locus or double-locus variants of the correct ST (i.e., where 6/7 or 5/7 of the MLST loci have exact matches, these are reported as STx-1LV or STx-2LV respectively by Kleborate). The Illumina-only and SUP+Illumina hybrid assemblies gave identical MLST results for all genomes. Compared to this gold-standard result, for ONT-only assemblies, the proportion of correct ST calls was 87.3% for SUP basecalled assemblies with or without polishing; 78.2% for HAC+Medaka; and 32.7% for non-polished HAC (see **Figure 2A**). Fast basecalled assemblies showed poor results (20% correct with polishing, 0 without). ONT-only assemblies based on SUP basecalled reads reliably identified lineage in all cases when allowing for single- or double-locus variant calls. Assemblies based on HAC basecalling also performed well, with 96.3% (HAC+Medaka) and 90.7% (HAC) matching the true ST within 0-2 locus variants, respectively. Interestingly, for isolate KSB1_10J, the SUP+Medaka assembly yielded two loci mismatches with the expected ST323, while the SUP assembly without polishing yielded a single mismatch (i.e., a more accurate result).

**Figure 2.**
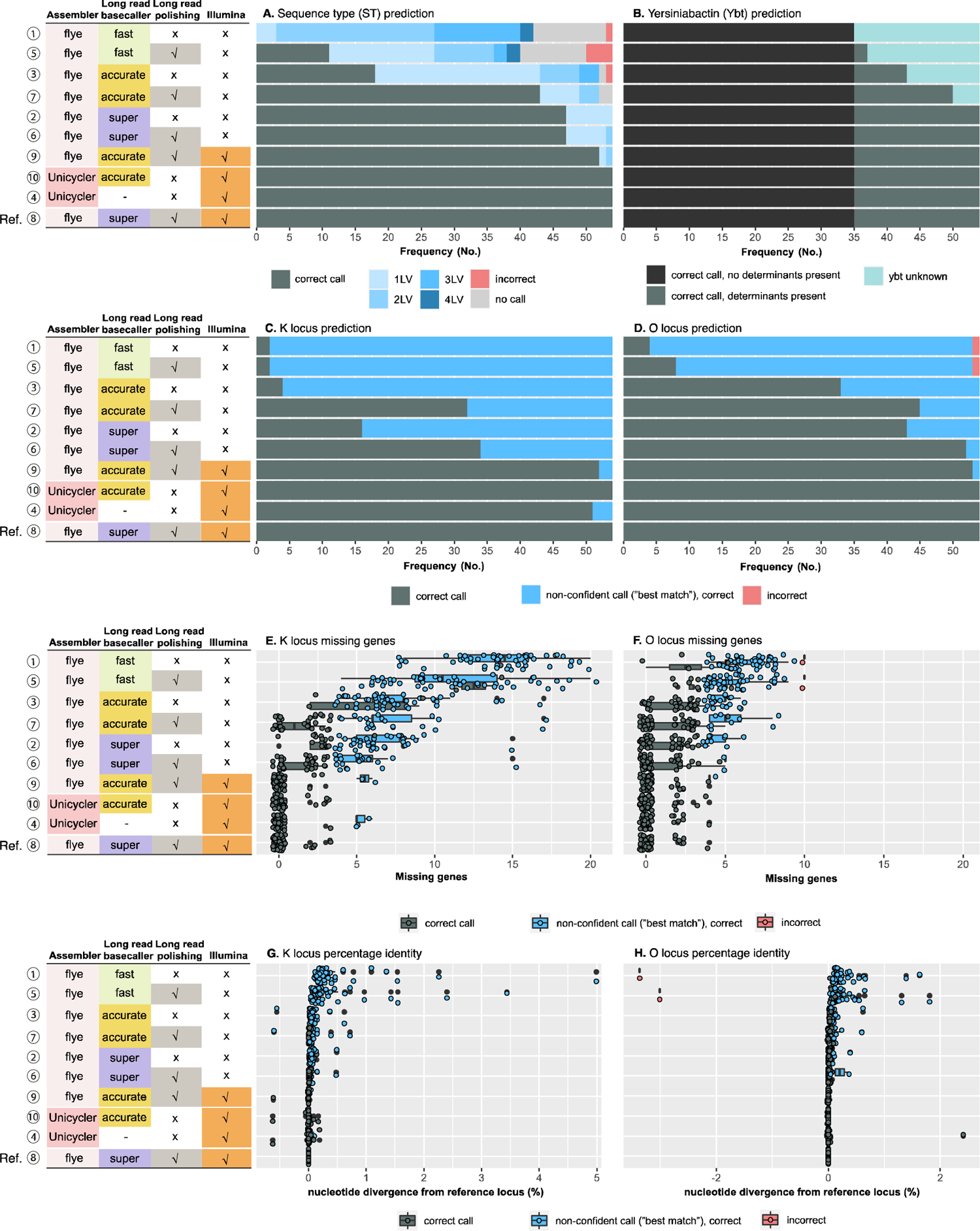
Genotyping accuracy for MLST, yersiniabactin and K/O loci using different assemblies. Each panel summarises accuracy of genotyping results across n=54 isolates, for each type of assembly (one per row as labelled on the left—numbered 1 to 10 as in Figure 1, and ordered based on overall performance), compared to the reference assembly (i.e., long-read first hybrid assembly using Flye and Medaka to assemble and polish with super-accurate basecalled reads, then polished with Illumina; last row per panel). **(A)** Multi-locus sequence type (ST) calling; 1LV, 2LV, 3LV, 4LV indicates the correct ST was predicted but the allele call was incorrect for 1, 2, 3, or 4 alleles. (**B**) Detection and typing of yersiniabactin; ‘correct call, no determinants present’ indicates the *ybt* locus was absent in the reference assembly and was (correctly) not detected in the test assembly; ‘correct call, determinants present’ indicates the *ybt* locus was present in the reference assembly and this was correctly identified and subtyped (to lineage level) in the test assembly; ‘determinants present, called as unknown’, indicates the *ybt* locus was present in the reference assembly, and this was correctly identified as present but could not be subtyped in the test assembly. (**C**) K locus and (**D**) O locus typing; ‘correct call’ indicates the correct locus was identified with confidence ranking of good or better; ‘non-confident call (“best-match”), correct’ indicates that the correct locus was identified but with low or no confidence. (**E**) and (**F**) show the number of open reading frames that are encoded in the reference K and O loci but were not detected in the test assemblies (labelled as ‘missing genes’ by the Kaptive genotyper, although in this case the nucleotide sequences are present but basecall errors result in disruption of the open reading frames). (**G**) and (**H**) show the nucleotide divergence between each assembly and the reference K and O loci, across the length of the detected loci.

Pathogenwatch calculates pairwise SNP distances (hereafter referred to as PW distances) across a set of 1,972 *K. pneumoniae* core genes and constructs a neighbour-joining tree based on these distances. **Figure 3** shows a PW-distance tree for all 540 assemblies analysed here. PW distances between ONT-only assemblies vs their corresponding hybrid reference ranged from 2–882 SNPs (median 17) for SUP+Medaka to 198–3390 SNPs (median 374) for unpolished Fast basecalled assemblies (**Figure 4**). Accordingly, the tree shows clear clustering of alternative assemblies for the same isolate, with SUP and HAC assemblies clustering closely with their corresponding hybrid references and Fast basecalled assemblies being distant relatives (see **Figures 3-4**, and interactive version of the tree in Microreact, [https://microreact.org/project/5chGLxaT1eVHKrThyc4b4J-nanopore-only-assemblies-for-genomic-surveillance-of-the-global-priority-drug-resistant-pathogen-klebsiella-pneumoniae]).

**Figure 3.**
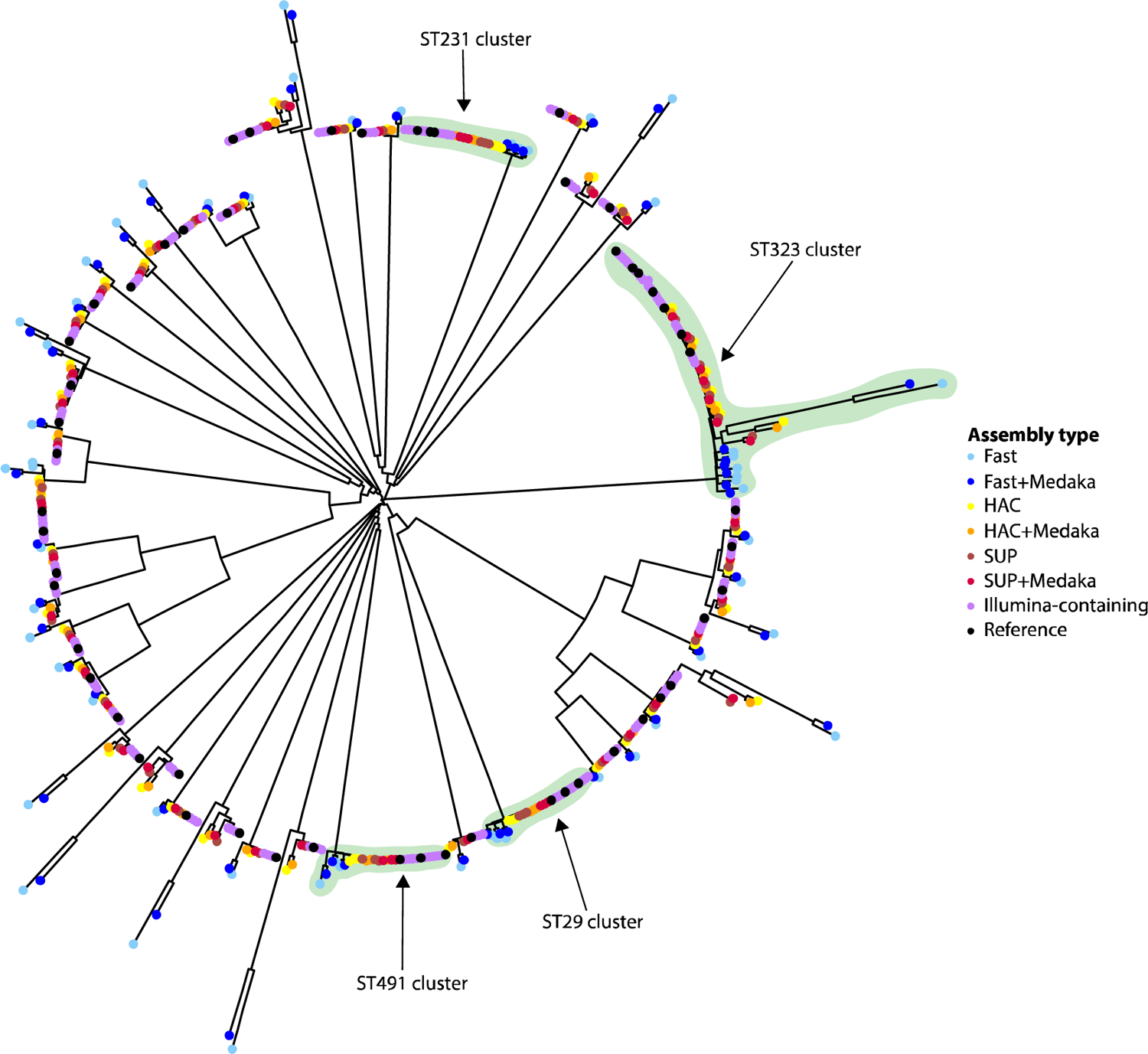
Neighbour-joining tree representing the phylogenetic relationships among the 540 genome assemblies, derived from n=54 *K. pneumoniae* isolates as outlined in Figure 1. The tree was constructed from the Pathogenwatch distance matrix (i.e. based on SNPs called in 1972 core genes) and is midpoint-rooted. Tips are coloured by assembly type, as indicated by the inset legend; note the reference assembly (i.e. long-read first hybrid assembly using Flye and Medaka to assemble and polish with super-accurate basecalled reads, then polished with Illumina) is coloured black and all other Illumina-containing genomes are grouped and coloured purple. Alternate assemblies of the same genome tended to cluster together but are non-zero branches because the sequences are non-identical due to basecalling errors. ONT/Illumina hybrid assemblies of the same isolated clustered more closely than ONT-only assemblies. Assemblies derived from the Fast model (±medaka, blue colours) consistently formed outliers separated by long branches, due to high rates of basecalling errors. Most of the study isolates belong to distinct *K. pneumoniae* lineages and are unrelated, except for sixteen isolates belonging to four transmission clusters (see Figure 6), these clades are highlighted in green and labelled by multi-locus sequence type (ST).

**Figure 4.**
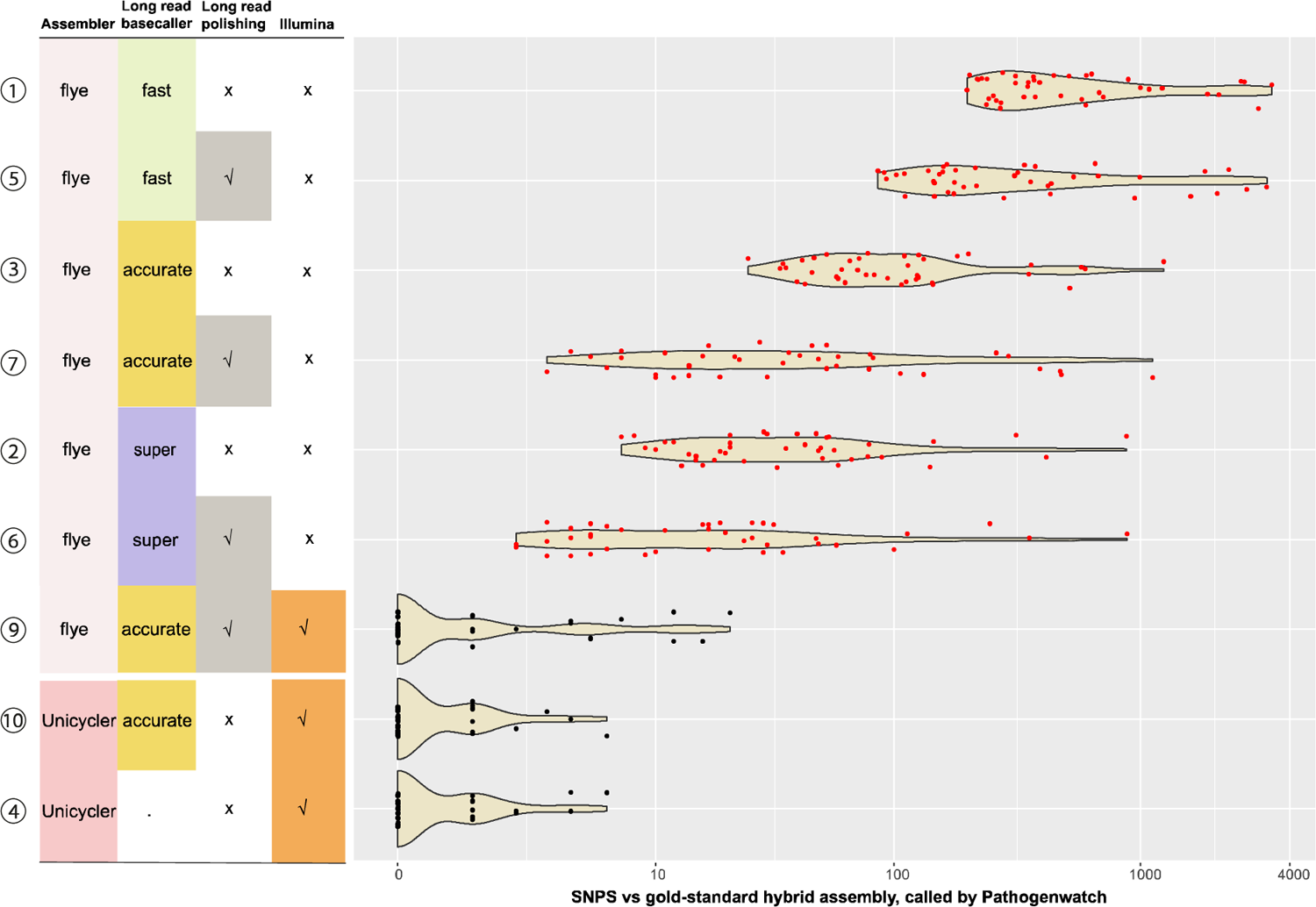
SNPs called vs reference assembly. A violin plot depicting the distribution of SNPs between each assembly vs the gold-standard hybrid reference assembly for the fifty-four study genomes based on a concatenated alignment of 1,972 genes (2,172,367bp) that make up the core gene library for *K. pneumoniae* in Pathogenwatch. Assembly types are numbered 1 to 10 as in Figure 1. Data points corresponding to ONT-only assemblies are coloured red. Within the cluster of ten assemblies for each sample, the SUP and HAC were consistently separated by a reasonable SNP threshold (≤10 SNPs); however, the Fast assemblies tended to form outliers (>100 SNPs). For clustering, only the SUP+Medaka assemblies reliably identified outbreak clusters (≤25 genome-wide SNPs threshold, which corresponds to ≤11 SNPs within the Pathogenwatch *K. pneumoniae* core genome typing scheme). HAC+Medaka polishing performed similarly to SUP without polishing, and polishing introduced errors into HAC- or Fast-basecalled assemblies.

We then constructed assembly method-specific neighbour-joining trees for the set of 54 isolates, using PW distance matrices constructed from either reference assemblies, or SUP, HAC or Fast basecalled and polished ONT-only assemblies. The trees based on SUP or HAC ONT-only assemblies yielded generally similar topologies to the reference tree (entanglement coefficients 0.31 and 0.35, Robinson-Foulds distances 20 and 22, respectively), but the Fast basecalled ONT-only tree was more divergent (entanglement coefficient 0.96, Robinson-Foulds distance 102; see **Figure 5** and **Supplementary Figures 3-4**). However, clustering by ST was evident in all trees regardless of basecalling and assembly method.

**Figure 5.**
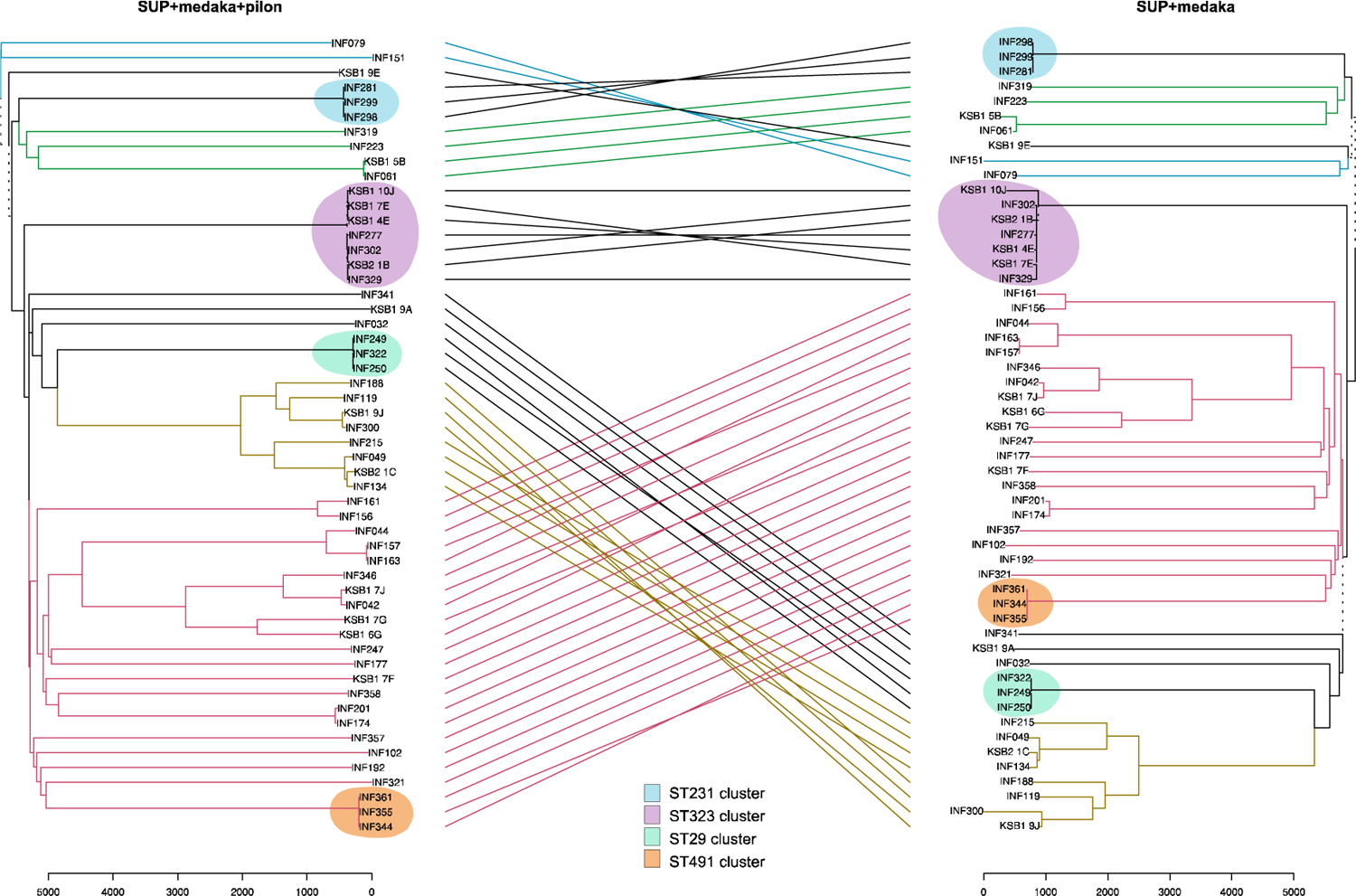
Comparison of trees constructed from ONT-only vs reference assemblies. Trees are midpoint-rooted neighbour-joining trees inferred from Pathogen pairwise distance matrices calculated from ONT-only (SUP+medaka) assemblies or ONT / Illumina long-read-first hybrid reference assemblies. Trees are shown as a tanglegram, which attempts to render the tree visualisations to maximise the alignment of tips, and links tips with lines (coloured to highlight shared subtrees), to facilitate comparison of topology between the two trees. The tanglegram yields an entanglement coefficient of 0.33 and a Robinson-Foulds distance of 20, indicating a good alignment. Coloured lines join matching tip labels for common subtrees, while grey lines indicate subtrees that are not common between the two trees. Discordant branch lengths on the ONT-only tree arise from basecalling errors, despite a good alignment.

Pairwise SNP distances can also be used directly, in the absence of trees, as a rule-in/rule-out indicator for potential transmission clusters. A consensus has emerged recently that a pairwise distance exceeding a threshold of 21–25 genome-wide SNPs is a reasonable basis for ruling out recent transmission of *K. pneumoniae* [65–67]. Based on comparisons using Illumina assemblies for n=270 unique *K. pneumoniae* clinical isolates, we estimate this equates to a PW distance of 10 SNPs (see **Methods** and **Supplementary Figure 1**). **Figure 6** shows PW distances calculated using reference assemblies, or SUP+Medaka ONT-only assemblies, for four groups of isolates in our data set that have previously been identified as part of nosocomial transmission clusters (based on a combined analysis of patient movement data and Illumina sequence, using genome-wide SNPs called from read mapping to an in-cluster reference sequence) [46, 87]. Clusters of ST29, ST231 and ST491 had pairwise PW distances 0-2 using reference genomes, but much higher PW distances using ONT-only assemblies (ranges 6-10, 4-4, and 6-20, respectively using SUP+Medaka; 13-23, 10-16, 19-28, respectively using SUP without polishing). Cluster ST323 was more diverse, with PW distances of 3-34 between reference genomes and 58-659 using ONT-only (excluding one outlier strain, KSB1_10J, the distances were 3-23 for reference assemblies, 58-118 for SUP+Medaka and 83-183 for SUP). Linear regression of polished ONT-only PW distances vs Illumina PW distances yielded a slope of ∼5 (adjusted r^2^=0.89, see **Figure 6**), suggesting a relaxed PW distance threshold of ∼50 could be suitable for calling transmission clusters using ONT-only (SUP+Medaka) data, although the relationship does seem to vary by ST (see **Figure 6**). We therefore conclude Pathogenwatch analyses of ONT-only assemblies are quite reliable for identification of lineage or ST (as described above), and thus could be reliably used to ‘rule out’ isolates of distinct lineages from suspected transmission clusters. However, more work is needed to establish suitable analytical methods for calculating and interpreting smaller distances within lineages, as is required to differentiate recent transmission chains from coincidental but independent infections with the same lineage.

**Figure 6.**
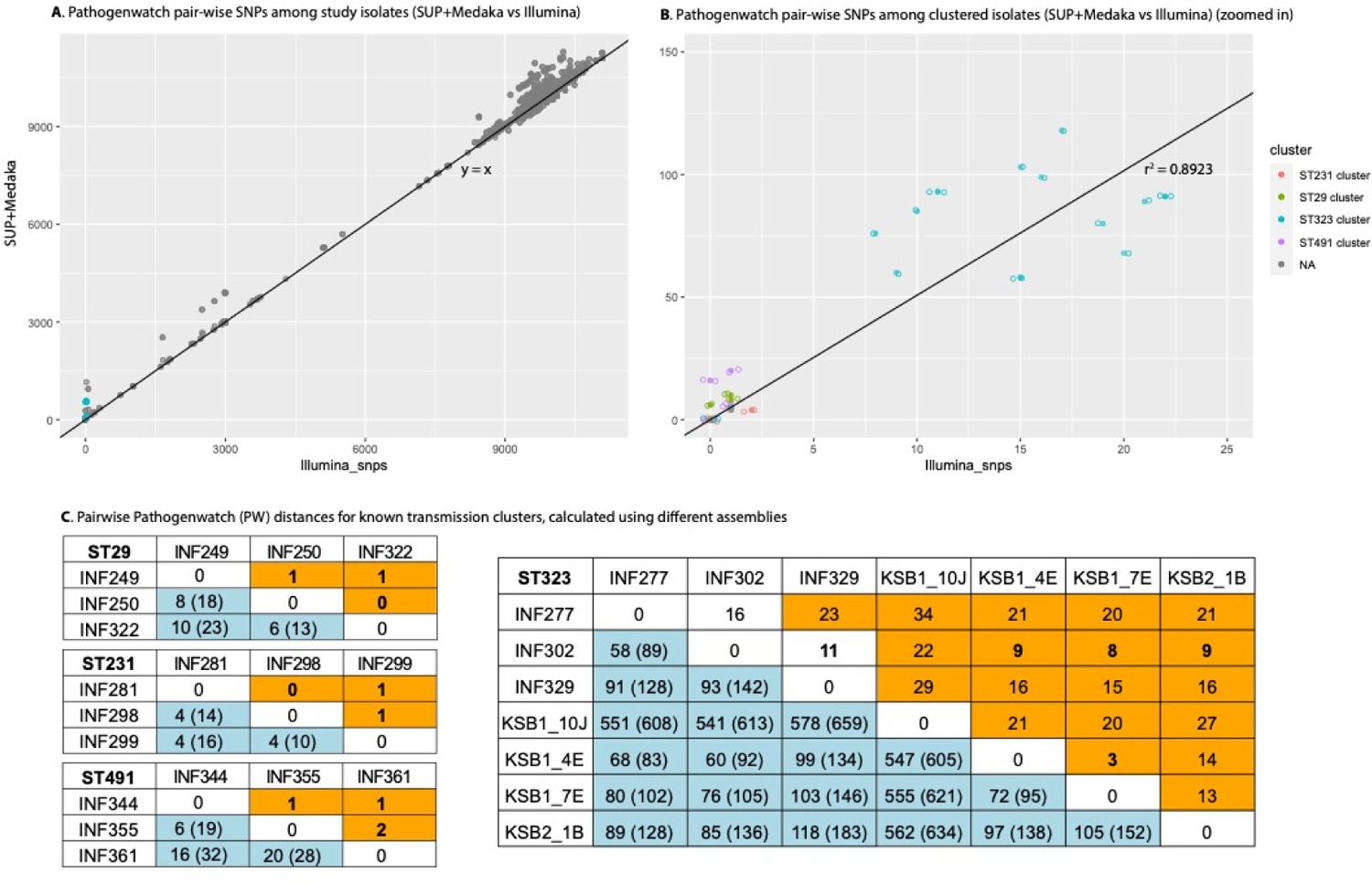
Comparison of pairwise Pathogenwatch distances between n=54 isolates, calculated from ONT-only vs Illumina assemblies. Y-axis (**A** and **B)** shows pairwise distances between isolates, calculated using SUP+Medaka assemblies for all isolates; x-axis shows pairwise SNP distances between isolates, calculated using Illumina assemblies for all isolates (based on the core gene set of 1,972 genes within the Pathogenwatch *Klebsiella pneumoniae* core genome scheme). Each data point represents a pair of isolates, coloured to indicate isolate pairs belonging to the same transmission cluster (as per inset legend). (**A**) All isolate pairs; line shows y=x. (**B**) Zoom in to transmission cluster isolates with ONT-based pairwise distances ≤150; linear regression line is shown (solid line), adjusted R^2^=0.8923 indicating very good model fit, slope=5.1. (**C**) Pairwise Pathogenwatch (PW) distances for known transmission clusters, calculated using different assemblies. PW distances calculated from reference assemblies are highlighted in orange; those calculated from ONT-only with SUP basecalling are highlighted in blue (numbers in brackets are those without Medaka polishing). Distances below the clustering threshold for PW distances, i.e., ≤11, are bolded.

### K and O locus typing

In *K. pneumoniae*, K and O locus types vary in their gene content, and typing via the Kaptive software is achieved by searching genomes for full-length reference K or O loci and confirming the presence of the expected genes within the best-matching locus [16, 17, 88]. K locus (KL) and O locus (OL) calls were in perfect agreement between Illumina and hybrid assemblies. Compared to these results as reference, all ONT-only assemblies correctly identified the best-matching KL type. However, the confidence reported by the Kaptive typing tool—which depends on the detection of intact genes (i.e., open reading frames) in the K locus and is thus expected to be impacted by basecall accuracy—varied widely for the different assemblies (see **Figure 2C**). The distribution of missing gene count, stratified by confidence, is shown in **Figure 2E**; this supports the expectation that the lack of confidence in KL calls is due to disruption of open reading frames in the locus, presumably due to basecall errors that result in frameshifts or stop codons in the expected coding genes. **Figure 2G** shows that nucleotide similarity (compared to KL reference sequences) along the K locus was quite high for all assemblies, with nucleotide divergence well below the Kaptive threshold for a confident call (≤5% divergence). This confirms that even small numbers of basecall errors are enough to disrupt open reading frames and thus reduce confidence in genotype calls.

The results were similar for O typing, although some incorrect OL calls were made with the Fast assemblies (1.85% each with Fast and Fast+Medaka; red colours in **Figure 2D**). A correct O1 prediction requires the detection of an extra gene (*wbbY*) outside the O locus [17, 89]; if this open reading frame is disrupted, O1 may be miscalled as O2. Unsurprisingly, the discrepancies with the Fast OL calls arose from O1 being erroneously reported as O2, due to frameshift mutations arising from basecalling/assembly errors affecting the unlinked *wbbY* gene.

### AMR determinants

Kleborate screens for acquired AMR genes by interrogating each input assembly against a species-aware modified version of the Comprehensive Antibiotic Resistance Database [45]. A total of 270 acquired genes were identified across the 54 reference genome assemblies (median 9 per genome, range 0-18). These acquired genes were mostly identified correctly from SUP+Medaka (95.8%), SUP (84.8%), HAC+Medaka (82.7%) and HAC (60.3%) assemblies, with lower recovery rates from Fast+Medaka (53.2%) and Fast (25.7%) assemblies. Kleborate also screens for mutations in chromosomal core genes known to be associated with AMR [45]. Here, reportable mutations were identified from reference assemblies in the *gyrA* and *parC* genes (substitutions associated with fluoroquinolone resistance, n=9 isolates), in *bla*SHV (substitutions associated with extended spectrum β-lactamase activity, n=27 genomes) and in *ompK35* and *ompK36* (truncations associated with reduced susceptibility to β-lactams, n=2 genomes). Truncations in *pmrB* and *mgrB* (associated with colistin resistance) are screened by Kleborate, but these genes were intact in all reference assemblies. The substitutions in *gyrA*, *parC* and *bla*SHV were recovered quite accurately from the SUP+Medaka (91.7%) and SUP (88.9%) assemblies, but less so using HAC+Medaka (83.3%), HAC (75%), Fast+Medaka (55.6%) and Fast (44.4%) assemblies (see **Figure 7a-b**). The two *omp* gene truncations were identified by all assembly methods, however, truncations of *omp*, *pmrB* and *mgrB* were also falsely identified in many cases (7.4% with SUP+Medaka, 7.4% SUP, 14.8% HAC+Medaka, 9.3% HAC, 77.8% Fast+Medaka, 27.8% Fast; see **Figure 7c-d**).

**Figure 7.**
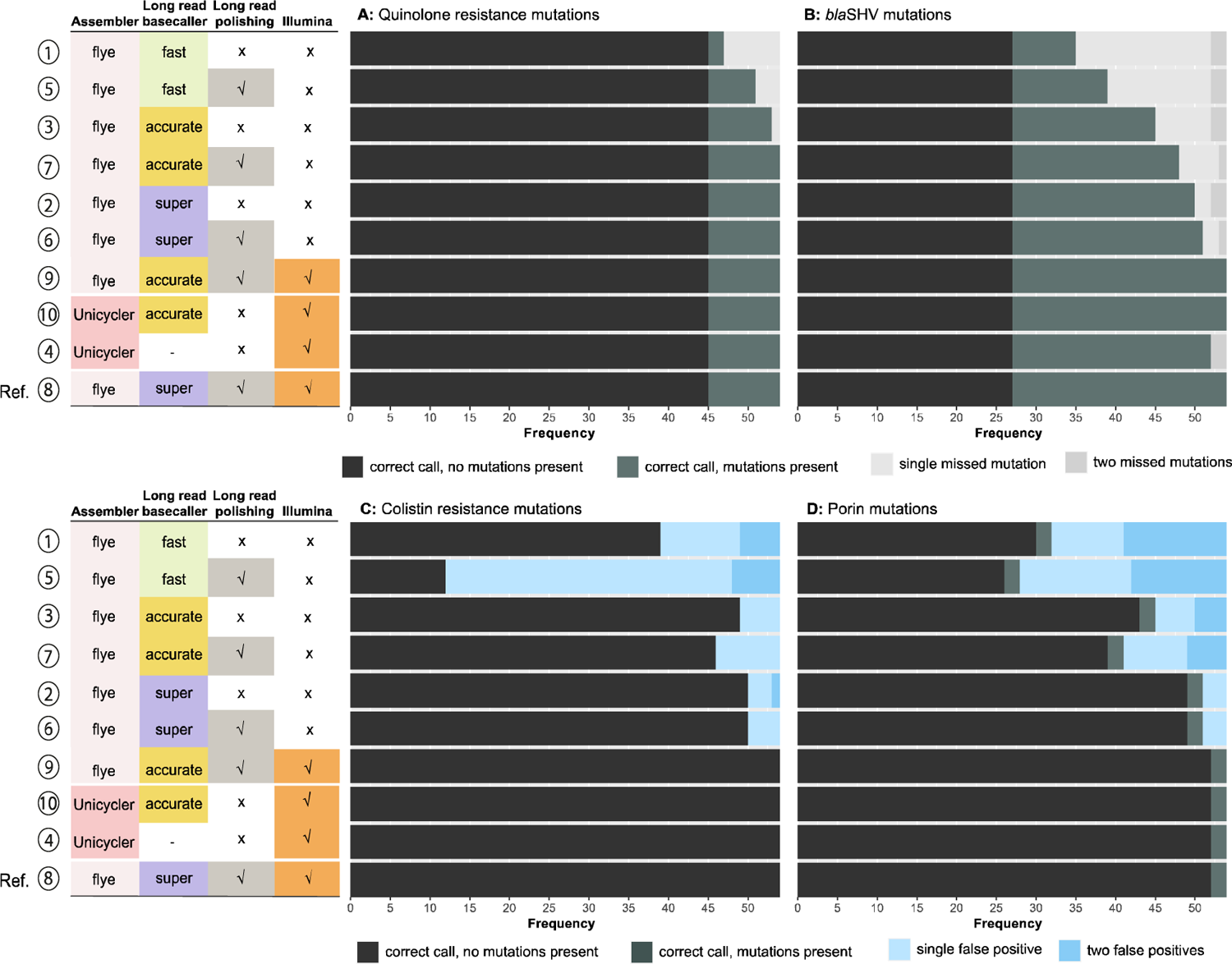
Detection of AMR-associated mutations in core chromosomal genes. Each panel summarises accuracy of genotyping results across n=54 isolates, for each type of assembly (one per row as labelled on the left—numbered 1 to 10 as in Figure 1, and ordered based on overall performance), compared to the reference assembly (i.e. long-read first hybrid assembly using Flye and Medaka to assemble and polish with super-accurate basecalled reads, then polished with Illumina; last row per panel). **(A)** Detection of fluoroquinolone resistance-associated mutations (GyrA-83I, ParC-80I, GyrA-83Y, GyrA-87N and GyrA-87Y). (**B**) Detection of mutations in *bla*SHV that are associated with a change in enzyme activity (35Q, 146V, 238S and 156D). (**C**) Detection of colistin resistance-associated mutations (disruption of MgrB or PmrB). (**D**) Detection of mutations in outer membrane porins associated with change in carbapenem susceptibility (truncation of OmpK35 or OmpK36, GD or TD insertions in OmpK36 loop 3).

As the presence of AMR determinants in genomes is often used to predict or explain resistance to clinically relevant drugs, we also considered the impact of ONT basecalling and assembly method on the task of identifying which isolates are likely non-susceptible to specific drug classes. **Table 1** shows the proportion of genomes that would be correctly classified as non-susceptible due to the detection of AMR determinants known to be present in the reference genome. SUP+Medaka assemblies performed well, with 88% – 100% of genomes with AMR determinants correctly identified as such across the various drug classes. HAC+Medaka also showed good detection rates (60% - 100%), while SUP results were more variable, ranging from 42-100% detection rate (see **Table 1**).

**Table 1:**
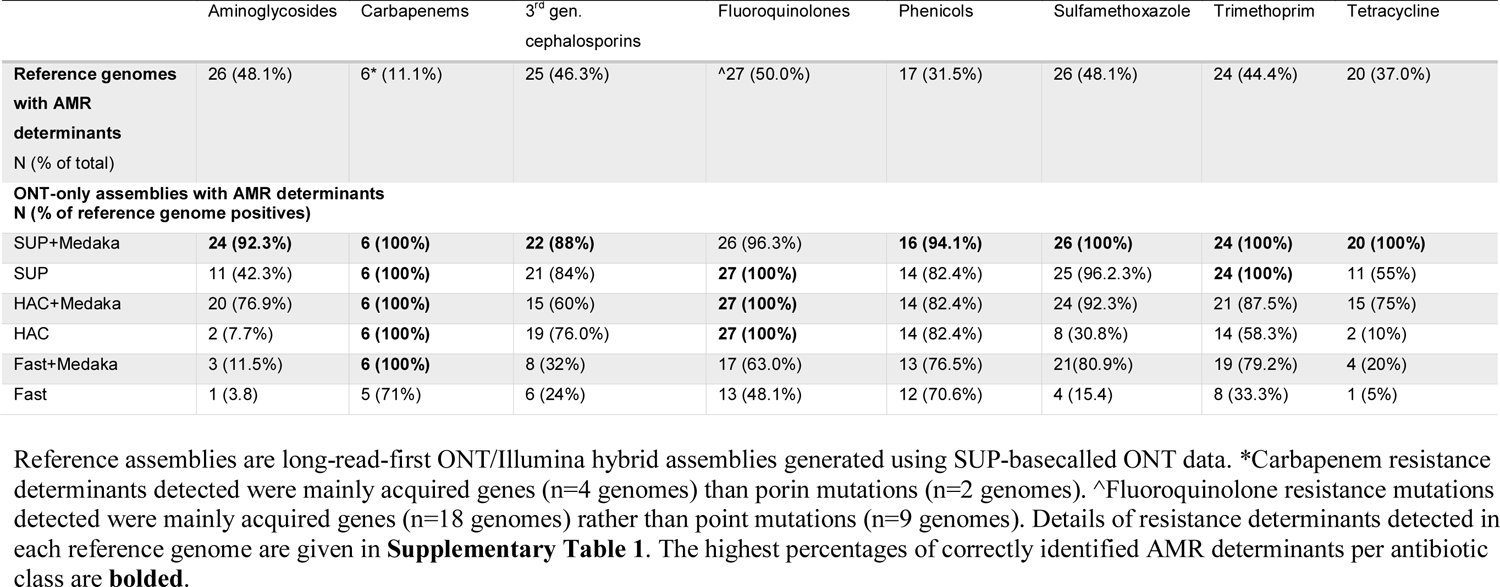
Accuracy of identifying non-susceptibility to specific drugs/classed based on detection of known AMR determinants

### Acquired virulence

Kleborate reports the presence of five major acquired virulence loci, including the ICE*Kp*-encoded siderophore yersiniabactin (*ybt*) and genotoxin colibactin (*clb*), plus three loci that are typically plasmid-borne and commonly linked with invasive infections caused by hypervirulent *K. pneumoniae*, namely, siderophores aerobactin (*iuc*) and salmochelin (*iro*), and the hypermucoidy locus *rmpADC.* Each of these loci comprises several genes, which are included in locus-specific MLST schemes for each [45, 90, 91]. Each locus will be reported, and an ST (or the closest match) assigned following the same logic to assign STs as for seven-gene MLST, if >50% of the genes contained in the locus are detected. Lineages of each virulence locus are inferred based on virulence STs, with each lineage pre-assigned to a set of virulence STs [90, 91]. Acquired virulence genes detected in our reference genomes include *ybt*, *clb,* and *iuc*. *Iuc* and *clb* were present in only two isolates each (*iuc* in INF079 and INF151, *clb* in INF079 and INF341), while *ybt* was detected in twenty isolates (36.4%). When present, the Illumina and hybrid assemblies correctly detected both *iuc* and *clb,* including correct lineage assignment. For the ONT-only assemblies, SUP (±Medaka) and HAC+Medaka correctly reported the *iuc* and *clb* STs and lineages, while the HAC and Fast (±Medaka) detected the loci as present but failed to identify a ST match and thus could not identify a lineage. Similarly, for *ybt*, the Illumina and hybrid assemblies correctly reported the presence or absence of the *ybt* locus, and accurately predicted the lineage when the locus was present (100% each). However, for ONT-only assemblies, only the SUP (±Medaka) matched the Illumina and hybrid in uniformly detecting *ybt* and assigning the correct lineage (**Figure 2B**). The other ONT-only assemblies all reported the presence or absence of *ybt* but failed to call the ST and thus to report a lineage (HAC+Medaka, 7.4%; HAC, 20.4%; Fast+Medaka, 31.5%, Fast, 35.2%). Notably, there were no false positives (i.e., reporting the locus present when absent), nor any false negatives (i.e., failing to detect presence of the locus when present).

### Limitations and future directions

An artefact of our sample collection is that there were few virulence plasmid-positive strains, so we could not assess other virulence genes apart from yersiniabactin. However, it is clear from the available results that genotyping based on detection of gene presence/absence is generally quite reliable using assemblies inferred from SUP-basecalled ONT reads, hence virulence gene detection is expected to be well tolerated generally. There were also very few strains in our collection with genuine AMR-associated truncations, so the call rate for these could not be accurately established. However, clearly, basecall errors frequently disrupt open reading frames, such that AMR-related truncations are subject to relatively high false-positive call rates. The same issue also affects the confidence of K and O locus typing: whilst the correct locus is almost always detected, the reported confidence of the call depends on the detection of open reading frames for all genes in the locus, which is rarely achieved using ONT data alone.

Our study considered *K. pneumoniae* only. However, the basecalling, assembly and genotyping tasks explored here are commonly applied across other bacterial pathogens. In particular, since *Enterobacterales* share common methylation patterns (which is a major determinant of basecalling accuracy) and AMR mechanisms, the results presented here can be considered informative as to the general accuracy of ONT-only analysis of the family more broadly.

Notably, laboratory and informatics methods for ONT sequencing are under constant development, e.g., new flowcells have recently been released with new (R10) pores; basecalling models can be trained on a larger or species-specific dataset, and assembly and polishing methods are under active development [92, 93]. These developments are expected to lead to improvements in the accuracy of ONT-only assemblies and therefore the accuracy of genotyping from such assemblies [36]. Therefore, our results should be considered a baseline from which improvements are expected to accrue. In addition, it is likely that the task of pairwise SNP calling could be more accurately performed using read-based variant calling approaches, rather than the assembly-based methods used here. However, there is currently a lack of SNP-calling tools designed specifically for bacterial ONT data (notably those trained on human data do not perform well on bacterial data due to differences in methylation [37]), so exploring this was beyond the scope of the current study which focused on the use of a consistent assembly-based approach to all analysis tasks (implemented in the freely accessible online Pathogenwatch platform).

Despite these limitations, our study provides a timely update on the level of accuracy that can be achieved with currently well-established and widely available protocols, including (i) R9.4 pores (which have been widely and stably used for several years now); (ii) out-of-the-box basecalling with Guppy, which can be done in real-time, and on-board with devices like Mk1C or GridION; (iii) a simple, rapid low-resource assembly with Flye (assessed in several benchmarking studies as the best singular assembler for ONT-only); and (iv) the well-established and stable Medaka polisher.

## Conclusions

Overall, our results show that MLST, K/O locus type, virulence and AMR determinants can be reliably identified from ONT-only genome assemblies. However, pairwise SNP distance estimation was less reliable and thus we propose that ONT-only analysis should be considered reliable only for rule-out of potential clusters using MLST-based lineage assignments, rather than being the sole trigger for specific actions that are usually based on a high suspicion of transmission (e.g., enhanced containment procedures in hospital settings), as it currently lacks the sensitivity needed for public health or infection control investigations. Where compute resources and/or time are limiting, our data indicate that compute resources are best directed towards basecalling (SUP model) over polishing.

## Author contributions

Conceptualisation: KEH. Methodology and Software: KEH, KLW, MMCL, RRW. Formal Analysis and Visualisation: EFN, HC, KLW, MMCL, KEH. Data curation: KEH, HC.

Supervision: KEH. Funding: KEH. Writing – Original Draft Preparation: EFN, KEH. All authors read and approved the final manuscript.

## Ethical statement

Ethical approval for the collection and sequencing of clinical isolates was granted by the Alfred Hospital Ethics Committee, Melbourne, Australia (Project numbers #550/12 and #526/13).

---

**Supplementary Figure 1.**
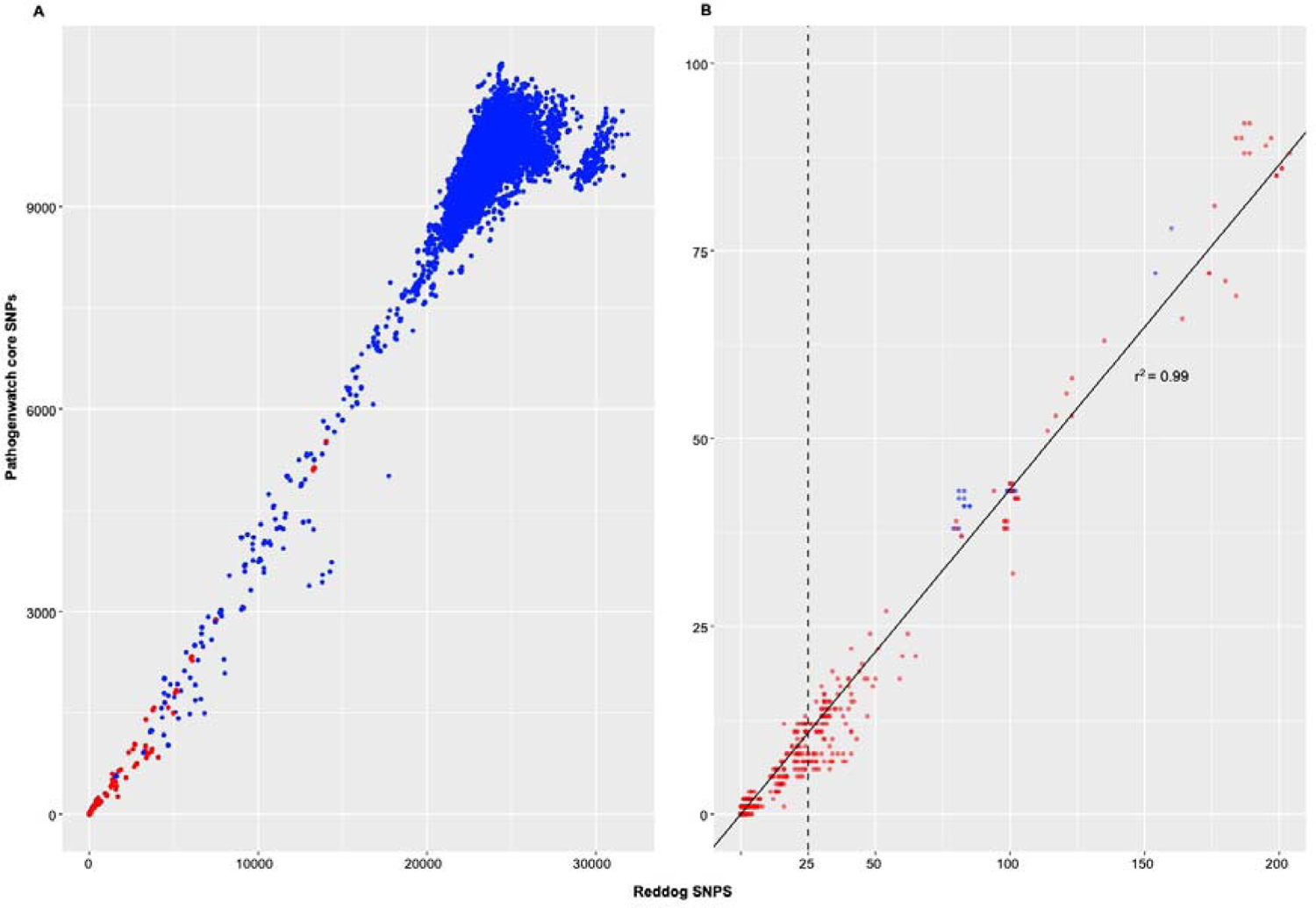
Comparison of Pathogenwatch distances with genome-wide mapping-based SNP distances, across n=270 diverse *Klebsiella pneumoniae* clinical isolates using Illumina data. Y-axis shows Pathogenwatch pairwise distances (based on the core gene set of 1,972 genes within the Pathogenwatch *Klebsiella pneumoniae* core genome scheme) using Illumina assemblies; x-axis shows genome-wide mapping-based pairwise SNP distances (based on all variant sites for which alleles were called in ≥95% genomes, when mapping Illumina reads to the NTUH-K2044 reference genome as described in reference 40). Data points represent pairwise distances, coloured to indicate pairs of genomes with the same (red) or different (blue) 7-locus sequence type. (A) All pairwise distances. (B) Zoom in to data points with pairwise genome-wide SNP distances ≤200; linear regression line is shown (solid line), adjusted R^2^=0.9914 indicating very good model fit, slope=0.432; dashed line shows x=25 genome-wide SNPs, which is a common threshold used to identify putative transmission clusters. According to the linear regression, this is equivalent to a Pathogenwatch distance of 10.8 SNPs (i.e., threshold n=10).

**Supplementary Figure 2.**
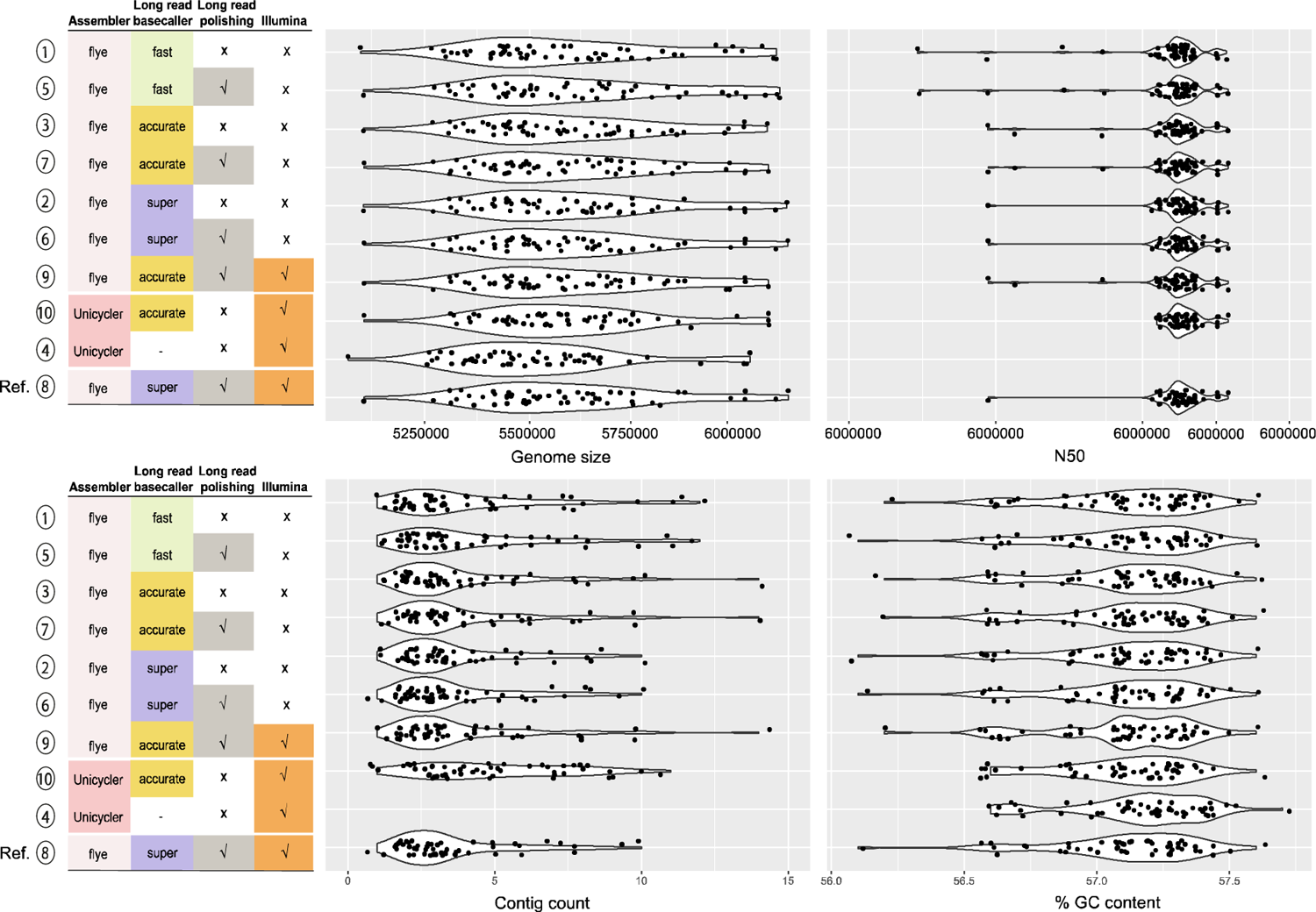
Distributions of sequencing metrics stratified by assembly type. Assembly types are numbered 1 to 10 as in Figure 1. (A) Genome size. (B) N50. N50 distribution for Illumina is not shown as it generates draft assemblies with N50 < 1 Mbp (range, 69194 – 612419; median, 274529) (C) Contig count. Similar to panel B, contig count distribution for Illumina is not shown as it generates draft assemblies with >20 contigs (range 32 – 315; median, 92) (D) %GC content. The y-axis shows the various assemblies under comparison, with the gold standard hybrid ONT / Illumina reference assembly (SUP+Medaka+pilon); while the x-axis displays the genome size, N50, contig count and %GC content distribution, respectively.

**Supplementary Figure 3.**
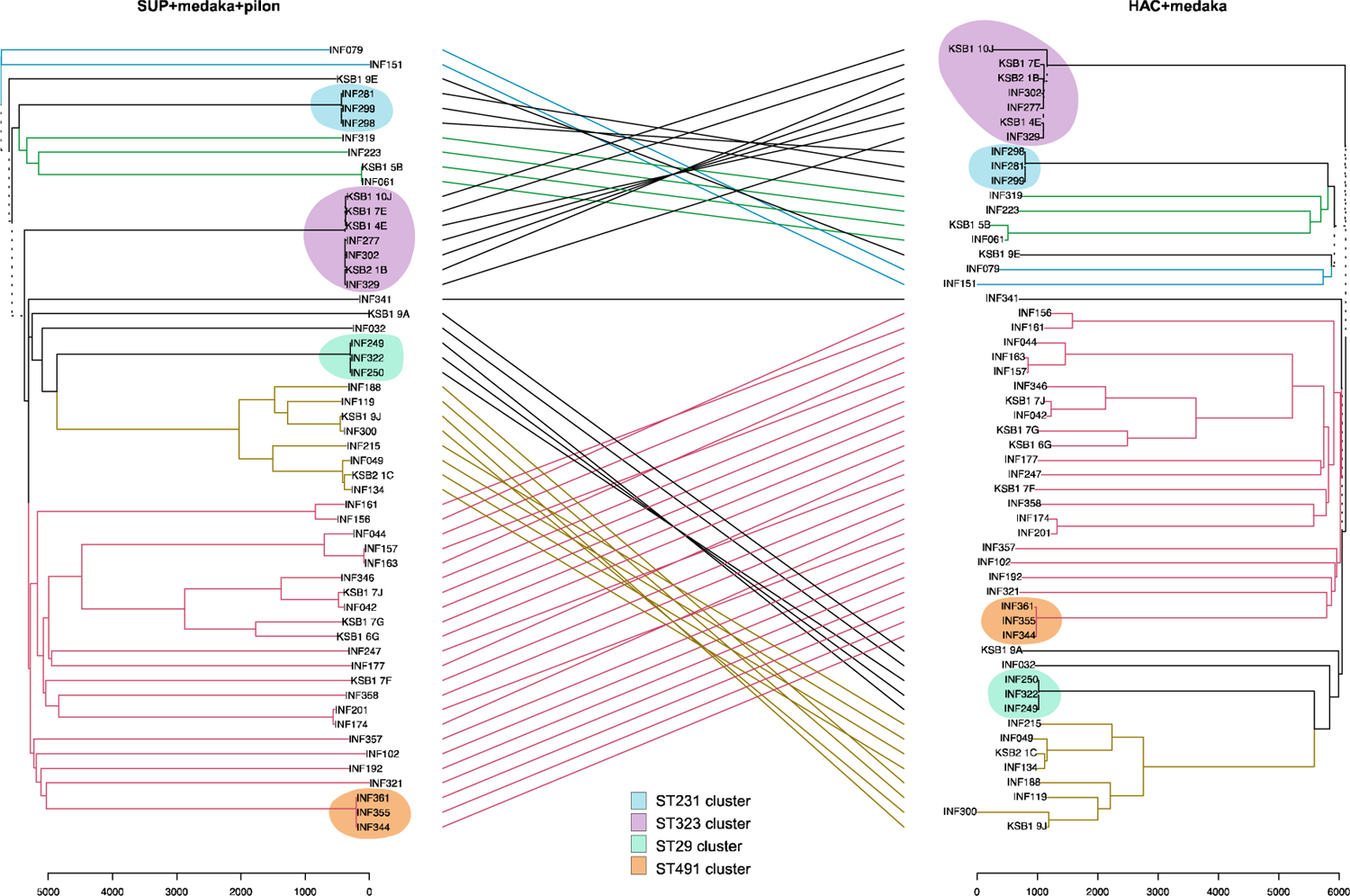
A tanglegram depicting the alignment between a Neighbor-Joining tree produced from HAC+medaka assemblies and the ONT / Illumina hybrid reference tree. The tree yields an entanglement coefficient of 0.35 and a Robinson-Foulds distance of 22, indicating a good alignment. However, several clades are discordant with the reference tree, represented by the dotted lines. Coloured lines join matching tip labels for common subtrees, while grey lines indicate subtrees that are not common between the two trees. Errors in the ONT-only assembly result in discordant branch lengths between the two trees.

**Supplementary Figure 4.**
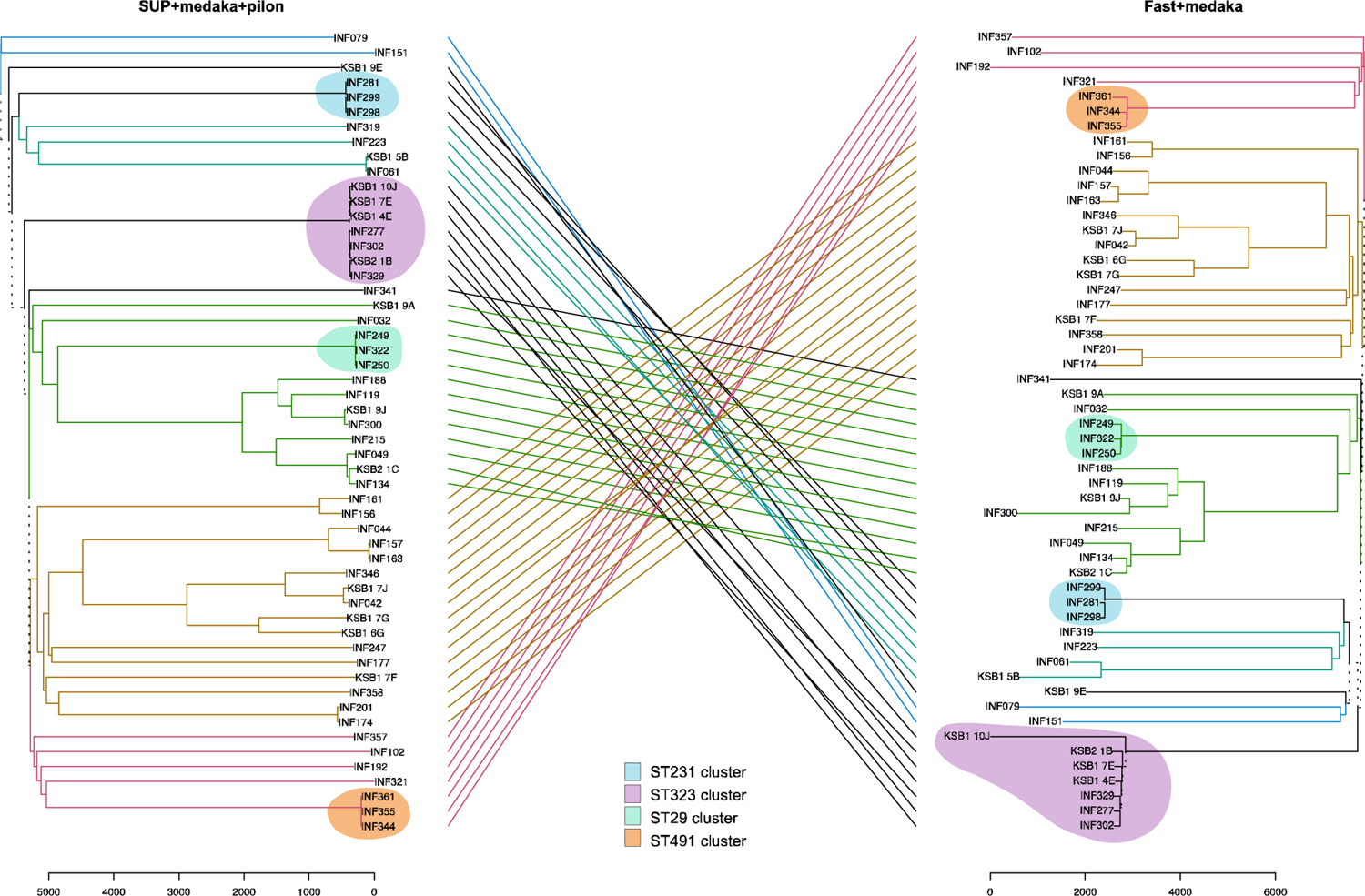
A tanglegram depicting the alignment between a Neighbor-Joining tree produced from Fast+medaka assemblies and the ONT / Illumina hybrid reference tree. The tree yields an entanglement coefficient of 0.96 and a Robinson-Foulds distance of 102, indicating a very poor alignment. Coloured lines join matching tip labels for common subtrees, while grey lines indicate subtrees that are not common between the two trees. Dotted lines represent distinct subtree present on either tree. Errors in the ONT-only assembly result in discordant branch lengths between the two trees.

